# Haemophilus, Antibiotic Therapy and the Airway Microbiome in Chronic Obstructive Pulmonarydisease

**DOI:** 10.1101/419127

**Authors:** Simon E Brill, Phillip L James, Leah Cuthbertson, Ana Zhu, Trevor Lawley, William OCM Cookson, Michael J Cox, Jadwiga A Wedzicha, Miriam F Moffatt

## Abstract

Chronic obstructive pulmonary disease (COPD) is a smoking-related illness affecting 64 million people worldwide. Airway infection drives recurrent exacerbations and lung function decline. Prophylactic antibiotics may prevent exacerbations but their use is a significant cause of population antimicrobial resistance.

We characterised the sputum microbiome by 16S rRNA gene analysis using 138 samples collected during a randomised controlled trial of prophylactic antibiotics in 71 patients with stable COPD. On comparing the profile of the microbiome obtained by sequencing to the isolates grown from samples using standard culture, there were similarities overall, although with a much narrower spectrum of genera on culture with under-representation of certain genera including *Veillonella* and *Prevotella.* There was concordance in the most abundant genera within samples and the number of isolates cultured reflected the measured bacterial diversity.

We found that at baseline the microbiota of 17 (24%) patients were dominated by *Haemophilus influenzae*, accompanied by narrowed microbial diversity and higher levels of sputum inflammatory cytokines. Different *H. influenzae* strains co-existed within individuals. Opportunistic whole genome sequencing of six *H. influenzae* isolates obtained during the study revealed that all were non-typeable *H. influenzae* (NTHI), with a range of different antibiotic resistance gene profiles, but an identical complement of virulence genes.

Administration of 13 weeks prophylaxis with moxifloxacin, azithromycin or doxycycline revealed distinctive changes in microbial communities for each group. *Haemophilus* numbers reduced by 90% compared to placebo only after moxifloxacin, and significant reduction in sputum cytokines occurred in patients dominated by *Haemophilus* at baseline. *Haemophilus influenzae* dominance defines COPD patients with active disease who may particularly benefit from antibiotics or vaccination.

## Introduction

COPD is a debilitating chronic lung disease characterized by episodes of symptomatic worsening associated with disease progression, healthcare utilisation and mortality (1-3). COPD is associated with structural and immune defects (4) and altered microbial communities within the airways. The presence of pathogenic airway bacteria is associated with exacerbation frequency and disease progression (5-6). Studies of the COPD microbiome with sequencing technology have so far used invasive sampling from small numbers of severely diseased patients (7, 8). There is an unmet need for larger studies relevant to the wider range of patients with COPD in the general population.

Long-term antibiotic therapy is one therapeutic option to prevent exacerbations (9). Long-term macrolide antibiotics may reduce the frequency (10, 11) and duration (11) of exacerbations and fluoroquinolones may be effective in selected patients (12). Macrolides also have potent anti-inflammatory effects (13), and there are no data to show their action on the microbiota in COPD. The airway bacterial load does not change with prophylactic antibiotic therapy (14), although there are significant increases in antimicrobial resistance in treated patients (10, 14) and the general population (15).

In order to understand better the effects of prophylactic antibiotic therapy and to streamline its use, we have investigated in detail the composition of the airway microbiome in COPD, using spontaneously expectorated sputum samples from a randomised controlled trial of prophylactic antibiotic therapy in patients with stable disease (14). In keeping with clinical practice, we have studied sputum samples as a surrogate for events in the lower airways. We have also carried out a direct comparison of the microbiome profile obtained using sequencing to that obtained using traditional bacterial culture, in order to help delineate the relationship of these newer techniques to those that are currently used to guide clinical practice. Furthermore, six isolates were available from stored samples from siz different patients, allowing us to confirm the species, antibiotic resistance gene profile and potential virulence by whole genome sequencing.

## RESULTS

The 71 patients who were sampled before and after treatment had moderately severe COPD, with a mean (SD) age of 69 (9) years, 52 (72%) male, mean (SD) forced expiratory volume in 1 second (FEV1) of 50 (14) % predicted and moderate airway obstruction (Table 1). We carried out quantitative polymerase chain reaction (qPCR) of the V4 region of the 16S rRNA and used identical primers to sequence the same region on the Illumina MiSeq platform. After quality control measures 138 of the original 142 samples were included for analysis, comprising a total of 9,113,146 sequence reads from 2259 distinct OTUs and 114 genera. The median sequencing depth was 61088 reads per sample (range 4,241 – 248,309), and samples were rarefied to a minimum sequencing depth of 4,241 reads unless otherwise specified.

**Table 1:** Patient characteristics at baseline*. *Plus–minus values are means ±SD; others are medians (IQR). FEV_1_: forced expiratory volume in 1 second; FVC: forced vital capacity; SGRQ: St George’s Respiratory Questionnaire; qPCR: quantitative polymerase chain reaction; IL: interleukin. Between-group differences were assessed using one-way ANOVA, Chi-squared test, or Kruskal-Wallis test as appropriate. There were no significant between-group differences at the significance level of 0.05 except for baseline FEV_1_ and FEV_1_ % predicted (p = 0.033 and p = 0.009, respectively).

|  | Combined | Treatment Group |  |  |  |
| --- | --- | --- | --- | --- | --- |
|  |  | Azithromycin | Doxycycline | Moxifloxacin | Placebo |
| N | **71** | 18 | 20 | 19 | 14 |
| Age - years | **69±9** | 68±9 | 71±7 | 71±8 | 67±10 |
| Male, n (%) | **52 (72)** | 12 (67) | 14 (70) | 14 (74) | 12 (86) |
| Current smoker, n (%) | **33 (46)** | 6 (33) | 9 (45) | 13 (68) | 5 (35) |
| Smoking history, pack-years | **49±32** | 50±24 | 40±29 | 55±29 | 53±48 |
| Exacerbation frequency in previous year, self-reported | **2 (1, 3)** | 2 (1, 3.75) | 2.25 (1, 4) | 3 (1, 3) | 1 (1, 2) |
| FEV<sub>1</sub>, litres | **1.4±0.6** | 1.1±0.5 | 1.5±0.5 | 1.5 ±0.5 | 1.7±0.7 |
| FEV<sub>1</sub>, % predicted | **49.5±14.5** | 39.7±16.1 | 52.5±13.1 | 52.8±11.2 | 53.5±13.9 |
| FEV<sub>1</sub>/FVC ratio | **0.5±0.1** | 0.4±0.1 | 0.5±0.1 | 0.5±0.1 | 0.5±0.1 |
| FVC, litres | **2.9±1.0** | 2.6±0.8 | 3.1±1.1 | 2.9±1.1 | 3.2±1.0 |
| SGRQ score | **49.1±15.7** | 47.9±17.6 | 47.4±15.3 | 53.1±12.6 | 47.7±18.3 |
| Total bacterial | **9.22±0.77** | 8.87±0.63 | 9.33±0.73 | 9.44±0.82 | 9.06±0.85 |
| load by qPCR,<br>log<sub>10</sub> 16S rRNA<br>gene copies/g<br>sputum |  |  |  |  |  |
| Haemophilus-<br>specific<br>bacterial load,<br>log<sub>10</sub> 16S rRNA<br>gene copies/g<br>sputum | **7.99±1.34** | 7.82±1.20 | 8.09±1.22 | 8.34±1.59 | 7.68±1.25 |
| IL-1β, log<sub>10</sub><br>pg/mL | **2.13±0.70** | 2.15±0.81 | 1.94±0.69 | 2.34±0.61 | 2.08±0.70 |
| IL-8, log<sub>10</sub><br>pg/mL | **4.03±0.78** | 4.04±0.74 | 3.89±0.87 | 4.27±0.73 | 3.85±0.77 |
| IL-6, log<sub>10</sub><br>pg/mL | **1.71±0.71** | 1.81±0.77 | 1.53±0.80 | 1.86±0.65 | 1.63±0.59 |

### Haemophilus dominance of the microbiome defines a distinct cluster of patients with higher airway inflammation and lower airway bacterial diversity

We examined the profile of the microbiome in these patients at baseline. Streptococcus was the most abundant genus in sputum samples taken before antibiotic therapy, accounting for 42% of all sequence reads, followed by *Haemophilus* (19%). Sequences from the genera *Veillonella, Actinomyces, Granulicatella* and *Prevotella* comprised 17% of sequence reads (Supplementary Table 1 and Supplementary Fig. 4).

We carried out an unsupervised hierarchical clustering analysis of OTU frequencies which identified clear subsets of patients, the largest of which (n=42) had *Streptococcus* as the proportionally most abundant genus by number of sequence reads (Fig. 1a). We observed a cluster of 17 (24%) patients in whom sequence reads from *Haemophilus* OTUs were most abundant. Two samples, also related to this cluster, were dominated by Moraxella. A single sample dominated by *Pseudomonas* clustered separately.

**Figure 1:**
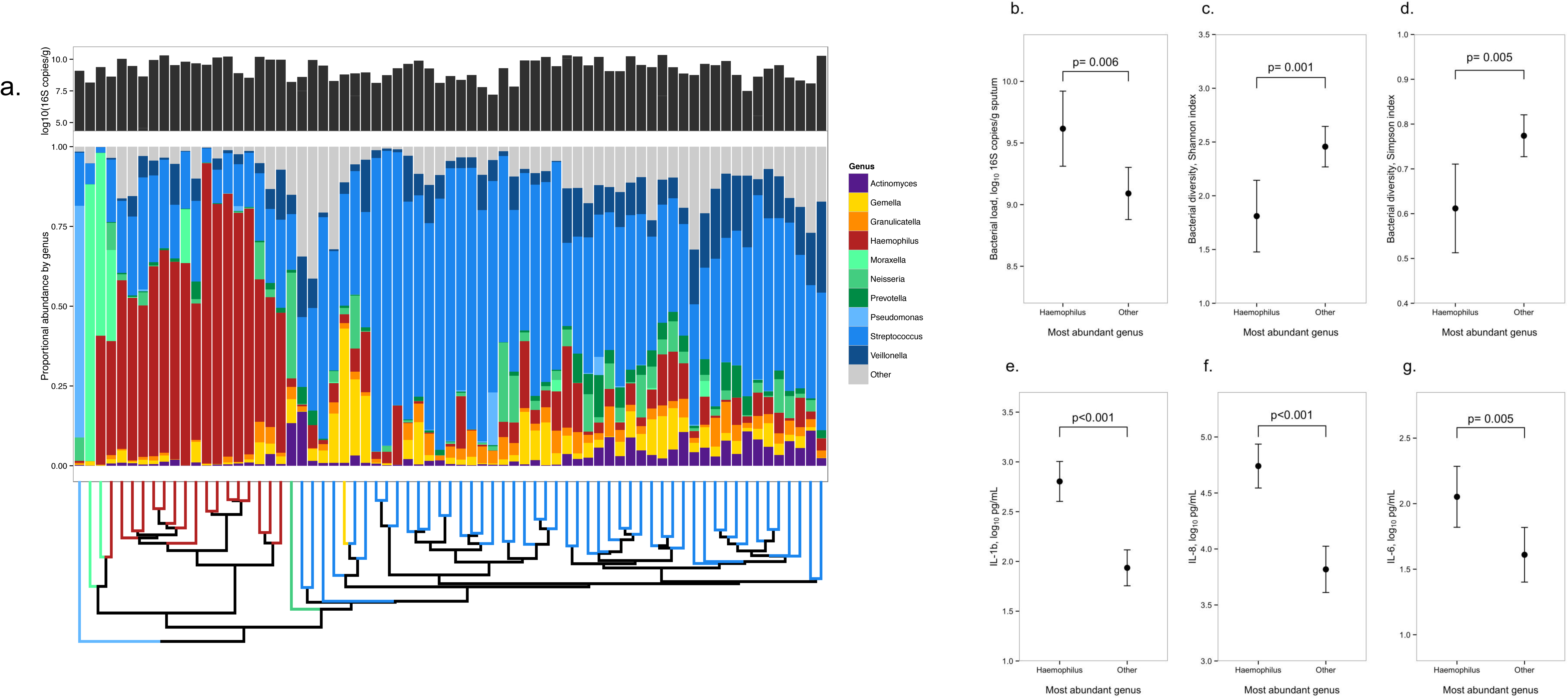
Profile of the COPD airway microbiome at baseline. **a**. Profile of the airway microbiome at baseline in 71 patients. The top panel shows the overall bacterial abundance by 16S rRNA gene qPCR within each sample. The central panel shows the proportional composition of the microbiome within each sample coloured by genus. Samples are clustered according to the unsupervised hierarchical clustering, the dendrogram for which below. This was constructed using the unweighted pair group method with arithmetic mean and the Bray-Curtis distance measure. The dendrogram leaf colours show the most abundant genus within that sample; the cluster of 17 patients dominated by *Haemophilus* species can clearly be seen. **b-g**: Significance of *Haemophilus* dominance in the microbiome at baseline. Bacterial load (b.), bacterial diversity (c. and d.), and inflammatory markers in sputum (e. – g.) according to whether the dominant genus in the microbiome at baseline was *Haemophilus* (n=17) or another genus (n = 54). Error bars show 95% confidence intervals around the mean. Unpaired t-tests were used for these comparisons. The patients in whom *Haemophilus* was the most abundant genus had higher levels of 16S rRNA gene copies, lower bacterial diversity, and higher levels of all three cytokines in sputum than others.

The mean (SD) bacterial load by 16S rRNA gene qPCR was 9.2 (0.8) log_10_ copies/gram of sputum. Sputum bacterial load was correlated with the inflammatory cytokine IL-1β (r=0.311, p = 0.009) but not FEV_1_, health status, exacerbation frequency, other cytokines, or inhaled corticosteroid use.

*Haemophilus influenzae* is a major recognized pathogen in patients with COPD (16). The cluster of 17 patients in whom *Haemophilus* was the most abundant genus had more 16S rRNA gene copies by qPCR (mean ± standard deviation 9.62±0.59 vs 9.09±0.78 log10 16S copies/g, p = 0.006, or 70% higher), Fig. 1b) than other subjects. In the *Haemophilus* dominated patients, we observed that the within-individual (alpha) bacterial diversity was significantly lower by the Shannon (1.81±0.65 vs 2.46±0.69, p = 0.001) and Simpson (0.61±0.19 vs 0.77±0.17, p = 0.004) indices (Fig. 1c & 1d). The levels of inflammatory cytokines were also notably higher (IL-1β [2.80±0.38 vs 1.94±0.65 log_10_pg/mL, p<0.001]; IL-6 [2.05±0.44 vs 1.61±0.75 log_10_pg/mL, p = 0.005]; IL-8 [4.74±0.37 vs 3.82±0.75 log_10_pg/mL, <0.001]) than non-*Haemophilus* dominant cases (Fig. 1e-g).

Correspondingly, there were strong correlations between the proportion of sequencing reads comprised by *Haemophilus* (proportional abundance) and the three inflammatory markers (IL-1b, r=0.56, p<0.001; IL-8, r=0.52, p < 0.001; IL-6, r=0.28, p = 0.02). We saw no significant differences in demographics, FEV_1_, previous exacerbation frequency or health status between groups (Supplementary Table 2).

### <u>The predominant Haemophilus</u> OTU within the COPD microbiome is *H. influenzae*

Of the 306 different *Haemophilus* OTUs present at baseline (1,208,824 sequence reads), 84% were from a single OTU (*Haemophilus*_1393, Fig. 2a). This was also the most abundant OTU in each of the *Haemophilus* dominated samples. Sequence alignment and phylogenetic analysis confirmed that this OTU clustered strongly within the branch containing *H. influenzae* and *H. haemolyticus* (Fig. 2b).

**Figure 2:**
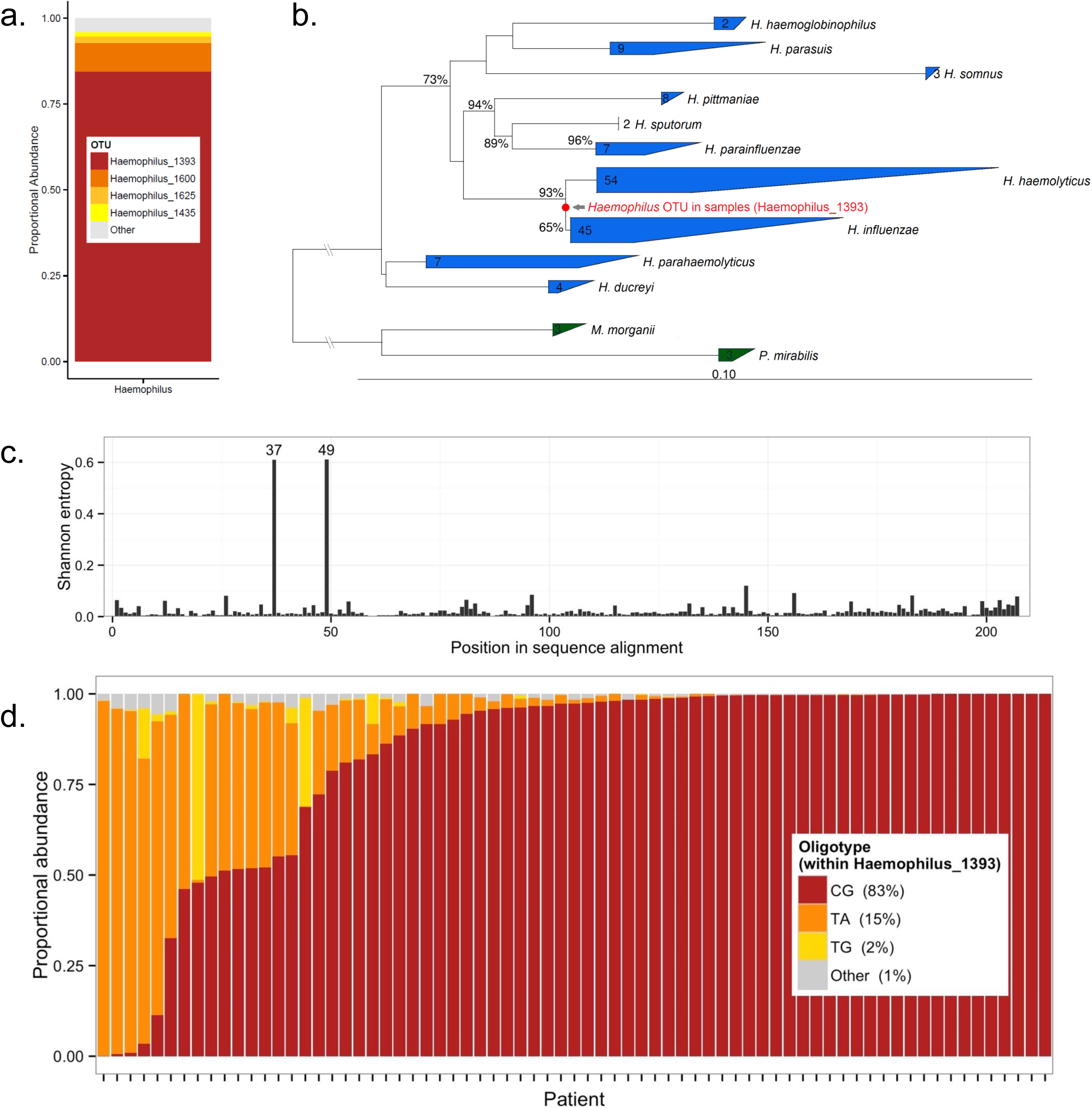
Characterizing *Haemophilus* species at baseline. **a**. Profile of individual *Haemophilus* operational taxonomic units (OTUs) in the microbiota of 71 patients at baseline. Eighty-four % is comprised of one OTU, assigned the label *Haemophilus*_1393. b. Phylogenetic tree showing *Haemophilus*_1393 as a close phylogenetic relation of *H. influenzae* and H. haemolyticus. The tree contains 141 representative sequences from other *Haemophilus* species and is rooted using the near neighbors *Proteus mirabilis* and *Morganella morganii*. The figures on each branch indicate confidences based on 1000 bootstrap replicates (values under 50% not shown). The figures within each leaf indicate the number of representative sequences contained therein. c. Shannon entropy resulting from oligotyping analysis of 1,019,560 sequences that were grouped within the OTU *Haemophilus*_1393. Entropy was analysed for each of the 206 positions within aligned sequences in order to detect information-rich positions. Two positions (37 and 49) display high entropy and were used to define distinct sub-clusters (‘oligotypes’) within this OTU. d. Proportional oligotype composition of the OTU *Haemophilus*_1393 within each patient at baseline.

Oligotyping is a supervised computational technique that allows the detection of meaningful sub-populations within closely related sequences, for example those assigned to a single OTU, and offers improved resolution over standard clustering techniques. Oligotyping of the 1,019,560 sequence reads within Haemophilus_1393 identified two base-pair positions of high entropy within the sequence alignment (Fig. 2c). Three distinct oligotypes, defined by the base differences C-G, T-A and T-G at these two positions, comprised 99% of the sequence reads (Fig. 2d). Notably, in approximately one quarter of patients there was more than one of these oligotypes present within the OTU. These are likely to represent different strains of the species.

In order to provide further confirmation of the species of this OTU, we examined the corresponding base-pair positions from the 16S rRNA gene sequences of 42 high-quality *H. influenzae* reference strains, finding C-G was present at these positions in 27 (64%), T-A in 13 (31%) and T-G in 2 (5%). Comparison of these positions to 96 other representative *Haemophilus* species confirmed that, while the C-G and T-G combinations were seen in other *Haemophilus* species, T-A was only present in reference strains of *H. influenzae*. Furthermore, *H. influenzae* was cultured from 9/17 (53%) of the samples where *Haemophilus* was the most abundant genus (compared to 3/54 [6%] where it was not) and was the only *Haemophilus* isolate from any of these samples. We were therefore confident that this OTU represents *H. influenzae*, comprises most of the *Haemophilus* present in the microbiome of our COPD patients, and that different strains of *Haemophilus* may co-exist within the microbiome.

### Whole genome sequencing of six *Haemophilus influenzae* isolates

Six *Haemophilus influenzae* isolates were archived during the study, coming from patients [ADD WHICH PATIENTS AND WHAT TREATMENT ARM THEY WERE IN]. These were whole genome sequenced, assembled and annotated. They were all identified as non-typeable *Haemophilus influenzae* owing to absence of both capsular regions I and II. Antibiotic resistance genes were annotated using the CARD database [REF] revealing that all six strains had at least one macrolide resistant gene, with a variable profile of additional resistances (figure 2e). The NTHI genomes were searched for a list 40 known virulence determinants [REF] revealing an identical set of 19 virulence genes to be present in all six isolates (Supplementary tables 1 and 2). This might suggest selection of particular NTHI strains adapted to the COPD lung. Two further analyses were therefore carried out to determine whether this was the case, a phylogenetic tree based on 40 universally conserved single-copy marker proteins and a pan-genome analysis of core (present in 90% of isolates in both these isolates and the Chiara et al 2014 genomes) and accessory genes. The six genomes did not cluster together by either method. A tanglegram comparison of a phylogeny built from a concatenation of the 19 virulence genes with the marker protein tree demonstrated agreement, indicating that selection does not appear to working at the strain level, though may still be occurring at the level of presence or absence of genes given the identical set of 19 virulence genes across the isolates (supplementary figure XX).

### Fluoroquinolone administration is associated with a particular decrease in *Haemophilus* taxa, while *Haemophilus*-dominated patients show an improved inflammatory response to antibiotic therapy

The patients were randomised to receive 13 weeks of treatment with azithromycin 250mg three times/week (18 patients), doxycycline 100mg daily (20 patients), moxifloxacin 400mg daily for 5 days every four weeks (18 patients) or 1 placebo capsule daily (14 patients). As previously reported (14), overall airway bacterial burden by 16S rRNA gene qPCR did not change significantly between the start and the end of treatment in any arm.

We examined whether treatment with antibiotics caused differences in overall community composition between treatment groups by performing permutational multivariate analysis of variance (PERMANOVA) at baseline and at the end of treatment. Prior to starting antibiotics we saw no significant effect of treatment allocation, but a treatment arm effect was present after the trial (R^2^ = 0.11, p<0.001) (Supplementary Fig. 6). There were no significant changes in within-sample diversity by the Shannon diversity index and number of observed OTUs in any treatment arm modeled against placebo.

Azithromycin treatment resulted in a decrease from baseline in eight non-*Haemophilus* OTUs, whilst seven OTUs (two from the genus *Haemophilus*) increased (Fig. 3a and Supplementary Fig. 7). Treatment with doxycycline resulted in a decrease in 29 OTUs, four of which were from the genus *Haemophilus*; this included a decrease of 85% in *H. influenzae* (Fig. 3b and Supplementary Fig. 7). The most profound changes were seen in the moxifloxacin treatment arm where a total of 107 OTUs decreased significantly, 80 (75%) of which were from the genus *Haemophilus*, including a decrease of 99% in H.influenzae. There was a corresponding increase in 37 OTUs, 15 (41%) of which were from the genus *Veillonella* (Fig. 3c, Supplementary Fig. 7). In the placebo arm, the only change was in one Pseudomonas OTU that increased significantly in abundance (Fig. 3d). These OTU-level changes were also seen in the overall profile of the microbiome after treatment, where there was near-total eradication of *Haemophilus* OTUs from the microbiome after moxifloxacin therapy but not following azithromycin or doxycycline (Supplementary Fig. 8).

**Figure 3:**
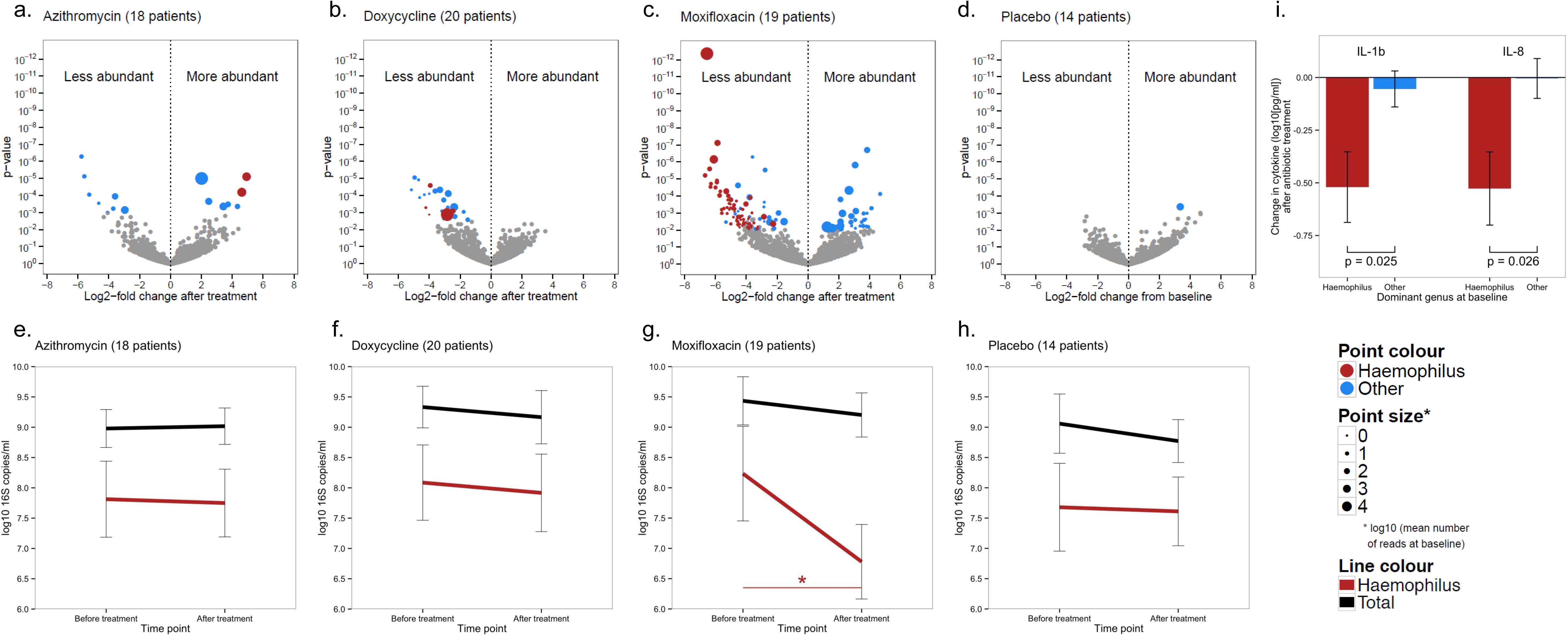
Changes in community composition and inflammatory cytokines following 13 weeks of antibiotic treatment with azithromycin (18 patients), doxycycline (20 patients), azithromycin (19 patients) or placebo (14 patients). **a-d**. Volcano plots displaying change in OTU abundance following treatment with azithromycin (a), doxycycline (b), moxifloxacin (c) or placebo (d). Each data point represents a single operational taxonomic unit (OTU). The OTUs highlighted (red: *Haemophilus*, blue: other genera) are those that changed significantly after controlling the false discovery rate at the 5% level. The size of the highlighted OTUs represents the log_10_ abundance prior to treatment start. There was a particular reduction in the abundance of *Haemophilus* OTUs following treatment with moxifloxacin. **e-h**. Changes in total bacterial load and *Haemophilus* load before and after antibiotic therapy in each treatment group. The bars represent 95% confidence intervals. Compared to placebo, the only significant change after treatment was the decrease in *Haemophilus* load following moxifloxacin treatment (g). * p = 0.02 when modelled against placebo in an adjusted analysis. **i.** Changes in the inflammatory cytokines IL-1β and IL-8 following antibiotic treatment (any) according to dominance of the microbiome at baseline (by *Haemophilus* [n=16] compared to other genera [n=41]) in all patients who received antibiotic therapy. Reduction in inflammation was significantly greater in those patients whose microbiota were *Haemophilus* dominated at baseline.

**Figure 4.**
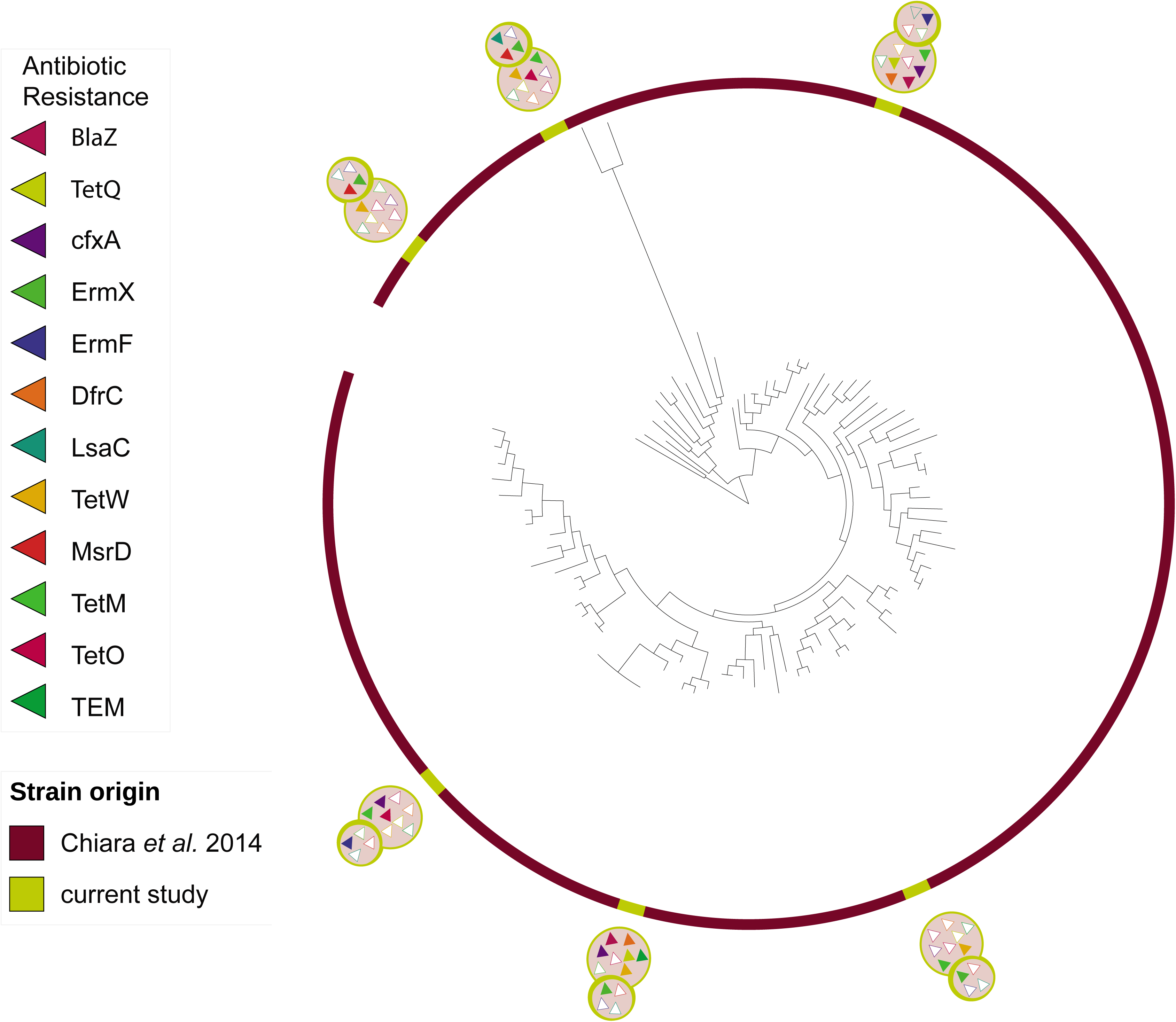
represents a phylogenetic tree based on concatenated alignment of 40 universal single-copy marker genes for 97 genomes (91 from Chiara *et al.* 2014 [1] and 6 specific to current study), using FastTree2 [2]. The antibiotic resistance genes for each of the 6 genomes are displayed above each sample in two superimposed circles. The antibiotic resistance annotation is based on CARD database [3]. Within the smaller circle are shown resistance to macrolide antibiotics [relevant to the study] and in the larger circle are shown resistance to other classes of antibiotics (beta-lactam, tetracycline and trimethoprim). Most of the genomes have at least a single resistance to macrolide.

Compared to placebo, there was a mean change in *Haemophilus*-specific 16S rRNA gene copies from baseline of 0.21 (95% CI -0.64 to 1.05; p = 0.62) log_10_ copies/gram after treatment with azithromycin, 0.20 (−0.60 to 1.01; p = 0.62) after doxycycline and -1.00 (−1.81 to -0.20; p = 0.02) after moxifloxacin. This corresponded to a 90% reduction in *Haemophilus* load with moxifloxacin therapy but not with azithromycin or doxycycline (Figs. 3e-f). Although levels of sputum cytokines did not change significantly following treatment in any arm, the change in *Haemophilus* load was overall predictive of the change in IL-1β (β-coefficient 0.13, 95% CI 0.05 to 0.22; p = 0.003) (Supplementary Fig. 9). Correspondingly, in the patients who received antibiotic treatment (n=57) there were greater falls in cytokine levels in the *Haemophilus* dominated group (n=16) than in the non-*Haemophilus* dominated group (IL-1β, -0.54±0.68 vs -0.05±0.65 log_10_pg/mL, p = 0.025; IL-8, -0.54±0.72 vs -0.03±0.67, p = 0.026; Fig. 3i). Patients whose microbiota are dominated by *Haemophilus* at baseline therefore showed a greater inflammatory response to antibiotic therapy.

## Discussion

We have shown that, in patients with stable COPD, dominance of the microbiome by the pathogen *H. influenzae* defines a group of patients who have more active disease with higher airway bacterial load and airway inflammation and lower bacterial diversity. Importantly, these patients also exhibited an improved inflammatory response to antibiotic therapy.

This expands on previous data linking bacterial colonization to exacerbation frequency (5) and disease progression (6). Defining ‘dominance’ of the microbiome by the most abundant genus provides a means of stratifying complex data and this has been shown in non-cystic fibrosis bronchiectasis to be clinically meaningful (17). We found agreement between the dominant genus and the most abundant isolate on culture, indicating that these DNA sequences originate from viable bacteria and that this dominance is clinically important. While studies published previously are too small to infer patterns individually, detailed profiles of the sputum microbiota of 92 clinically stable COPD patients have been reported across six studies (18-23). Twenty-eight (30%) of these were dominated by the genus *Haemophilus* or its parent phylum *Proteobacteria* (Supplementary Table 4). This is entirely consistent with our results.

Expectorated sputum was used during this study and carries a risk of contamination by oral microflora. Previous studies that have attempted to avoid this by using invasive specimens obtained bronchoscopically (7) or from lung explants (8) are necessarily small and lacking statistical power. The ubiquitous use of sputum microbiology in clinical practice indicates that minor contamination does not prevent expectorated sputum from providing a valid means of assessing the thoracic airway microbiome.

The quarter of patients in whom *H. influenzae* dominated the microbiota exhibited much higher levels of inflammatory cytokines and bacterial load than other cases. This is consistent with previous reports (24) using species-specific techniques but has not previously been shown in the full context of the airway microbiome. Airway inflammation predicts exacerbation frequency (25) and lung function decline (26), and these patients are therefore at high risk.

*Haemophilus* abundance was also associated with a narrowed diversity of species within the airway, and when the burden of *Haemophilus* reduced following moxifloxacin treatment there was a corresponding increase in the abundance of other bacterial genera. That there was no corresponding increase in measured diversity within the microbiota of the moxifloxacin-treated patients may reflect that the expanding taxa were from genera already present at baseline. These dynamic changes suggest that *Haemophilus* is able to dominate the microbiome, leaving a vacant ecological niche when removed. The fluoroquinolone moxifloxacin exhibited particularly potent activity against *Haemophilus* species. While this is in line with previous reports (27), the near-total eradication of *Haemophilus* from the airway microbiome has not previously been demonstrated. Oligotyping was successfully used in order to improve the 16S rRNA gene sequencing identification of *Haemophilus* spp. OTUs as *Haemophilus influenzae*, despite short sequences being available. It also revealed co-infection with multiple strains of *H. influenzae*, which may have implications for antibiotic therapy.

The relatively minor effect of azithromycin on the airway microbiome is surprising, particularly as it has efficacy against *Haemophilus* (28). Macrolides may therefore prevent exacerbations by attenuating bacterial outgrowth following viral infection (29, 30), or via unmeasured effects on bacterial function. Alternatively, macrolides have a wide spectrum of immunomodulatory effects (13) that may underlie the reduction in exacerbation frequency. The dose used here however is lower than that used in subsequently published trials (10) and this may have resulted in a reduced effect on airway bacterial communities. Although doxycycline also had a lesser effect than moxifloxacin it did result in the attenuation of several OTUs including *H. influenzae*.

Long-term antibiotic therapy is not without risk, particularly the induction of antimicrobial resistance (14, 15). We identified at least one macrolide resistance gene in each of six isolates that were whole genome sequenced, though this does not necessarily translate to *in vivo* resistance. A better understanding of antibiotic effects on the airway microbiome will allow therapy to be tailored to those who will benefit most. Importantly, vaccination against non-typeable *Haemophilus influenzae* may spare antibiotic use in selected patients (16).

A restricted set of 19 virulence genes of the 40 screened for was found in each of the six isolates, all of which were from six different COPD patients. This suggests that there may be some selection of particular virulence strategies in NTHI in COPD, many of these genes are involved in adhesion. However, this is small opportunistic set of genomes and systematic sampling and sequencing of a much larger number of NTHI isolates from COPD patients is required to confirm or refute this.

Molecular characterisation of whole sputum communities allowed us to define *Haemophilus* dominance in our patients in a manner that was not possible with classical culture. Our 16S analyses have given novel insights into the response of all the airway bacteria to antibiotics, and we suggest that community analyses should underpin future therapeutic trials of antimicrobial therapy in COPD.

## METHODS

### Participants and Study Design

Patients were included from a previous single-blind randomised controlled trial of antibiotic therapy in moderate-severe COPD (clinicaltrials.gov registration NCT01398072) for whom stored sputum was available for analysis (14). Patients were over 45 years old with stable COPD (forced expiratory volume in 1 second [FEV_1_] <80% predicted and FEV1/forced vital capacity ratio <0.7) and symptomatic chronic bronchitis. Patients were randomised to receive either azithromycin 250mg three times/week, moxifloxacin 400mg daily for 5 days every four weeks, doxycycline 100mg daily or 1 placebo capsule daily. Clinical data and spontaneously expectorated sputum were collected at baseline and at treatment end 13 weeks later. Ethical approval was obtained from King’s College Regional Ethics Committee (reference 11/LO/0932). All patients signed written informed consent prior to study entry.

### Laboratory Analysis of Sputum

Quantitative culture was performed at the time of sputum collection (31). The remaining sputum was homogenized and aliquots stored at -80°C pending batched further analysis. Sputum plugs were selected to minimize the risk of oral sample contamination. The sputum cytokines interleukin (IL)-6, IL-8 and IL-1β were analyzed using high-sensitivity enzyme-linked immunosorbent assay kits (RD Systems, Abingdon, UK). Lower detection limits were 0.70, 3.5 and <1.0 pg/mL respectively. DNA extraction was carried out using the FastDNA Spin Kit for Soil (MP Biomedicals, Santa Ana, USA). Triplicate quantitative polymerase chain reaction (qPCR) assays for the bacterial 16S ribosomal RNA (rRNA) gene were performed using a ViiA 7 real-time PCR system running ViiA7 Software Base v1.1 (Life Technologies, Paisley, UK). 16S rRNA gene copy numbers per gram of sputum were extrapolated from the cycle threshold for each sample and corrected for dilution.

Extracted purified DNA from samples was barcoded and amplified using quadruplicate PCR. The dual-barcoding approach was used (32) with 16S rRNA gene primers S-D-Bact-0564-a-S-15 AYTGGGYDTAAAGNG and S-D-Bact-0785-b-A-18 TACNVGGGTATCTAATCC (33), targeting the V4 region of the 16S rRNA gene, and Nextera v2.0 barcodes (Illumina, Chesterford, UK). Sequencing was carried out using the Illumina MiSeq platform (Illumina, Chesterford, UK). Both 16S rRNA gene qPCR and sequencing used identical primers targeting the same region of the 16S rRNA gene.

Demultiplexing, quality trimming, clustering of sequences into distinct operational taxonomic units (OTUs) and representative set picking were performed using Quantitative Insights into Microbial Ecology version 1.9.0 (34). Representative sequences were aligned against the Silva 115 NR database clustered at 97% similarity. Taxonomy was assigned using the Ribosomal Database Project naive Bayesian classifier (35) and an OTU table was constructed for analysis. Prior to analysis, negative controls were examined for known laboratory contaminants (36) and these reads were filtered from the dataset. Samples with <1000 reads remaining were removed. Further detail on the above steps, and the rationale behind the antibiotic treatments, is provided in the Supplementary Data. The dataset for this study has been uploaded to the European Nucleotide Archive, accession number PRJEB11966.

### Statistical Analysis

Read numbers were rarefied (randomly subsampled) to a uniform sequencing depth equal to that of the minimum retained sample after quality control, 4,241 reads. Summary variables were expressed as mean ± standard deviation (SD) or median (interquartile range) and log-transformed as appropriate. Hierarchical clustering was performed using the unweighted pair group method with arithmetic mean and the Bray-Curtis measure of between-group (beta) diversity. The relative biomass of *Haemophilus* within a sample was estimated by multiplying the qPCR bacterial load by the proportion of the sequencing reads that were from the genus *Haemophilus*. Continuous data were compared using Pearson’s correlation coefficient and significant variables used to fit a multiple regression model against IL-1β levels at baseline. Changes in *Haemophilus* load against placebo were assessed using a multivariate regression model adjusted for baseline *Haemophilus* load and baseline % predicted FEV_1_. Phylogenetic tree construction and sequence comparisons used ARB version 6.0.1 (www.arb-home.de) with the SILVA SSU reference database version 119.1 (www.arb-silva.de). Oligotyping (37) was carried out using version 2.0 of the pipeline (www.merenlab.org).

Differential OTU counts before and after treatment were modelled using the negative binomial distribution in the DESeq2 package (38) version 1.6.3 and the false discovery rate controlled using the Benjamini-Hochberg correction (39). Statistical analysis was performed using R statistics version 3.1.3 (www.r-project.org) and the phyloseq package (40) version 1.10.0.

### Genome sequencing of isolates

Details of this can be found in supplementary materials. Briefly, 6 available isolates of *Haemophilus influenzae* had DNA extracted and submitted for sequencing at the Wellcome Trust Sanger Institute, genomes were assembled and annotated for virulence genes, antibiotic resistance cassettes and phylogenetic trees constructed to compare these isolates to other non-typeable *H. influenzae*.

## SUPPLEMENTARY STATEMENTS

### Acknowledgements

The authors would like to thank James Allinson at Imperial College for assistance in collecting the study data, Claire Eckold, Sarah Thurston and Timothy McHugh at University College London for assistance with carrying out the quantitative culture, Raymond Sapsford at Imperial College London for help with processing samples, and Steven Cowman at Imperial College London for advice regarding Fig. 1. The authors would also like to acknowledge and thank all the patients who participated in this study.

### Author contributions

JAW and SEB contributed to the original study design; SEB, PLJ, MJC, WOC, JAW and MFM conceived and planned the further work presented here. SEB collected samples and study data, processed samples, and performed the cytokine analyses. SEB, LC and PLJ performed the 16S qPCR and gene sequencing. SEB and PLJ performed the sequence processing, quality control analysis and oligotyping. SEB, PLJ and MJC constructed the phylogenetic tree. SEB performed the initial data analysis and wrote the first draft of the paper. All authors contributed to data interpretation and further analysis. All authors were involved at all stages of the editing process and have read and approved the final manuscript.

